# Can human embryonic stem cell-derived stromal cells serve a starting material for myoblasts?

**DOI:** 10.1101/119693

**Authors:** Yu Ando, Marie Saito, Masakazu Machida, Chikako Noro, Masataka Takahashi, Masashi Toyoda, Akihiro Umezawa

## Abstract

A large number of myocytes is necessary to treat intractable muscular disorders such as Duchenne muscular dystrophy with cell-based therapies. However, starting materials for cellular therapy products such as myoblasts, marrow stromal cells, menstrual blood-derived cells and placenta-derived cells have a limited lifespan and cease to proliferate *in vitro*. From the viewpoints of manufacturing and quality control, cells with a long lifespan are more suitable as a starting material. In this study, we generated stromal cells for future myoblast therapy from a working cell bank of human embryonic stem cells (ESCs). The ESC-derived CD105^+^ cells with extensive *in vitro* proliferation capability exhibited myogenesis and genetic stability *in vitro*. These results imply that ESC-derived CD105^+^ cells are another cell source for myoblasts in cell-based therapy for patients with genetic muscular disorders. Since ESCs are immortal, mesenchymal stromal cells generated from ESCs can be manufactured at a large scale in one lot for pharmaceutical purposes.

## INTRODUCTION

Duchenne muscular dystrophy is an intractable genetic disorder and effective therapies have not yet been developed. Novel approaches to treat Duchenne muscular dystrophy include small molecules, gene therapy, biologics such as cytokines and cell-based therapy (Asakura, 2014; Miyagoe-Suzuki and Takeda, 2010). Among these advanced therapeutic approaches, regenerative therapies have been focused due to recent advances of pluripotent stem cells with different types of reprogramming technologies (Higuchi et al., 2015; Santostefano et al., 2015). *In vitro* expansion of quality-controlled stem cells and transplantation into patients with degenerative diseases in an allogeneic manner can be one of the ideal therapeutic scenarios. Somatic cells such as myoblasts, marrow stromal cells, menstrual blood-derived cells and placenta (amnion, cholate plate, umbilical cord)-derived cells have been introduced as starting materials for cellular therapy products (Cui et al., 2011; Cui et al., 2007; Hida et al., 2008; Kawamichi et al., 2010; Toyoda et al., 2007). However, these somatic cells have a limited lifespan and cease to proliferate *in vitro*, and thus sufficient numbers of cells cannot be prepared to treat muscles of a whole body in cell-based therapies. From this viewpoint, cells with a long lifespan are more suitable for starting materials.

Human pluripotent stem cells such as embryonic stem cells (ESCs) and induced pluripotent stem cells (iPSCs) are immortal, and can therefore be a good source of large number of cellular therapy products with one lot for genetic muscular disorders (Asakura, 2014). In addition to immortality, ESCs and iPSCs exhibit pluripotency, i.e. capability to differentiate theoretically into almost all types of cells including myoblasts and their progenitor cells (Barberi et al., 2005). As a therapeutic cellular products, myoblasts and mesenchymal stromal cells are considered most suitable. In this study, we generated mesenchymal stromal cells from ESCs for the production of cellular therapy products to treat patients with genetic muscular disorders (Assereto et al., 2016; De Paepe et al., 2016). We developed a novel protocol to manufacture mesenchymal stromal cells from ESCs with certified materials that had been analyzed for viruses.

## RESULTS

### Generation of mesenchymal stromal cells

To generate mesenchymal stromal cells from human ESCs, we propagated sees2 cells on mouse embryonic fibroblasts (MEFs) and formed embryoid bodies (EBs) for 4 days on a feeder layer of freshly plated gamma-irradiated mouse embryonic fibroblasts (Figure 1). The EBs were then transferred to the collagen-coated flasks and cultivated for 60 to 70 days. The upper adherent cell layer was detached to obtain a resource of mesenchymal stromal cells.

**Figure 1.**
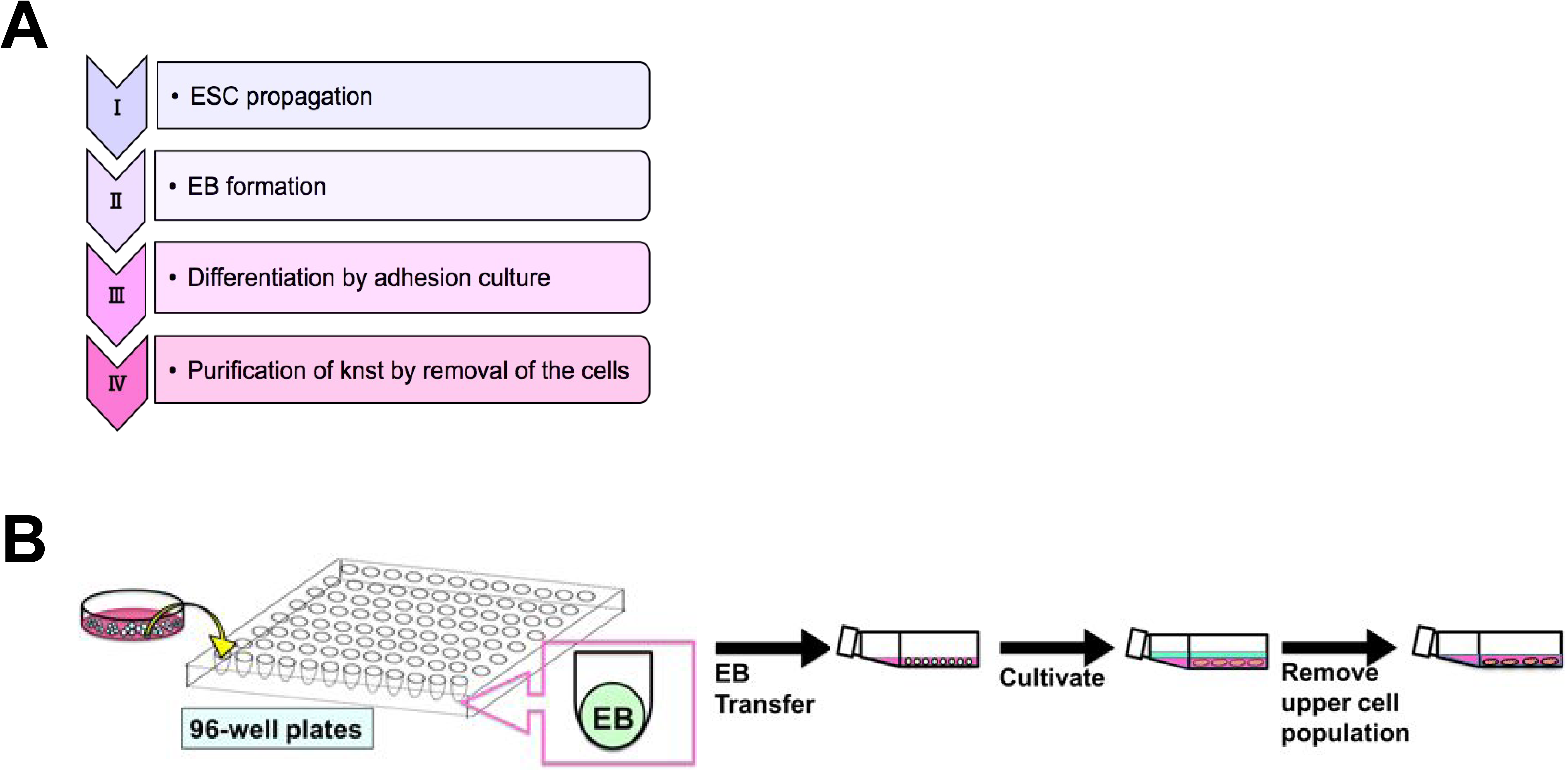
Generation of mesenchymal stromal cells from sees2. A. Step-by-step manufacturing process. B. Scheme for generation of mesenchymal stromal cells from sees2.

### Propagation of mesenchymal stromal cells

We repeated generation of mesenchymal stromal cells from sees2 cells in 4 different independent experiments (#3, #14, #23, #25), and investigated proliferation rate of the mesenchymal stromal cells (#3, #14, #23, #25) for over 50 days (Figure 2A). The mesenchymal stromal cells rapidly proliferated in culture and propagated continuously, however stopped replicating, became broad and flat, and exhibited SA-β-galactosidase activity as indicated by blue staining of their cytoplasm at passage 11 (Figure 2B, C). The enlargement of the cell size was passage-dependent.

**Figure 2.**
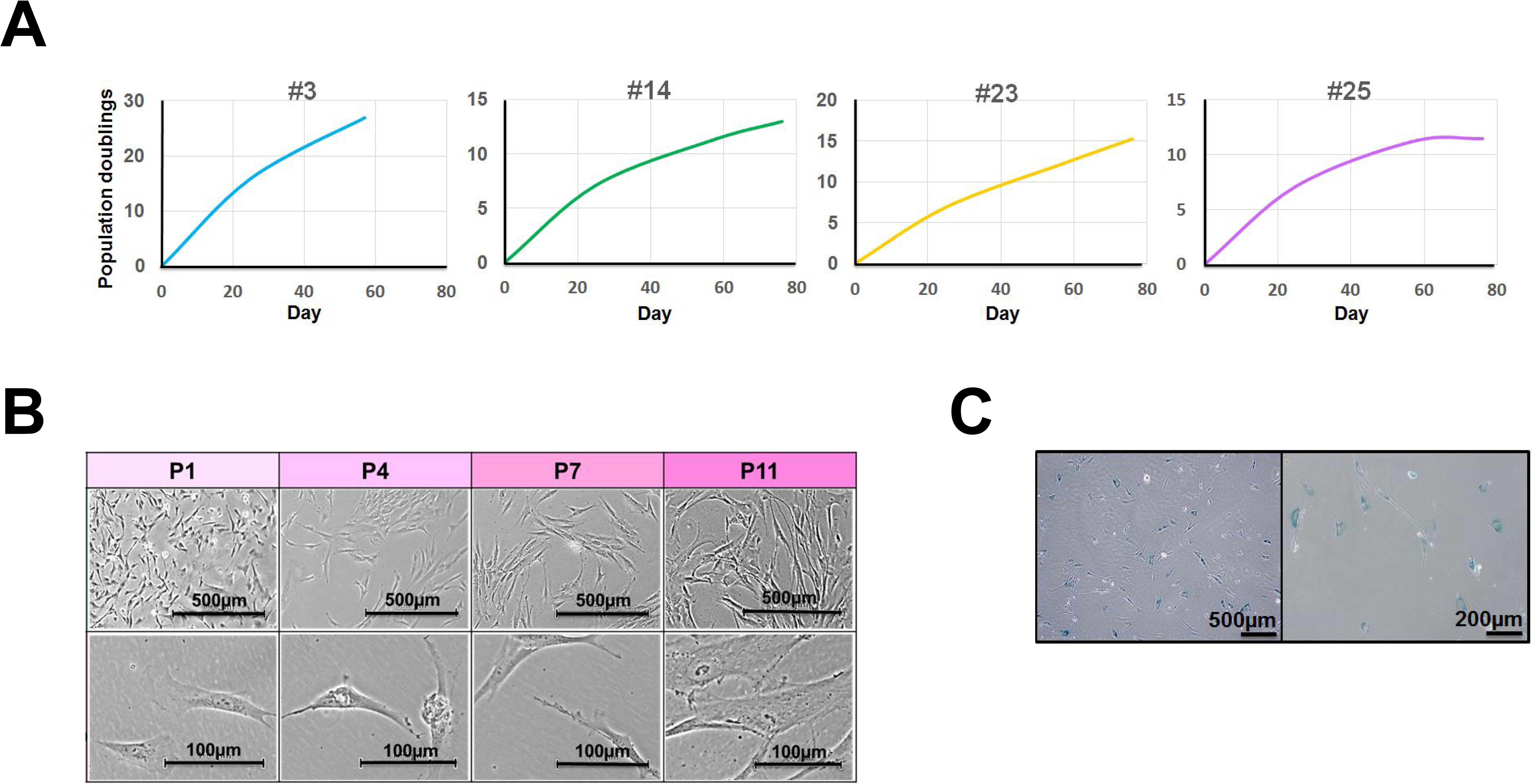
Characterization of mesenchymal stromal cells. A. Growth curve of mesenchymal stromal cells (knst#3, #14, #23, #25) B. Phase contrast photomicrography of mesenchymal stromal cells (knst#2: passage 1, 4, 7, and 11) C. Senescence-associated beta-galactosidase stain (knst#3, passage 11).

### Flow cytometric and karyotypic analysis

Flow cytometric analysis revealed that the mesenchymal stromal cells #2 and #3 were positive for CD90, CD105 and HLA-ABC, and negative for HLA-DR (Figure 3A). The expression level and pattern of these markers remained unchanged after 3 or 4 passages (12 or 16 population doublings). Karyotypic analyses of the mesenchymal stromal cells #2 and #3 were performed at passage 3 and 2, respectively (Figure 3B). They were found to be diploid and not to exhibit any significant abnormalities. The chromosome number of both #2 and #3 was 46 without exception.

**Figure 3.**
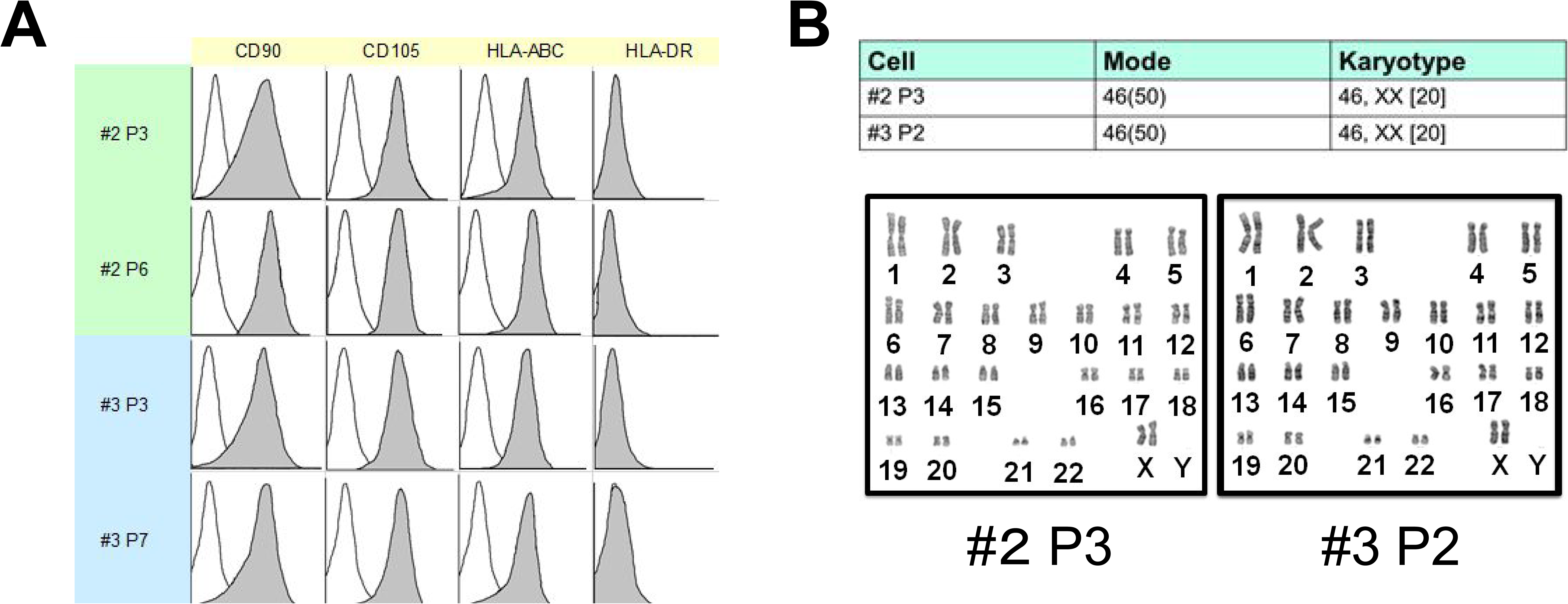
Flow cytometric analysis and karyotypic analysis. A. Flow cytometric analysis of knst#2 (passage 3 and 6) and knst#3 (passage 3 and 7). B. Karyotypic analysis of knst#2 (passage 3) and knst#3 (passage 2).

### Global outlook by hierarchical clustering and principal component analysis (PCA)

To investigate myogenic potential, mesenchymal stromal cells were analyzed, depending on gene expression levels. Hierarchical clustering analysis based on all probes, mesenchyme-associated genes, and stem cell-associated genes revealed that the mesenchymal stromal cells were categorized into the same group in a passage-dependent manner (Figure 4A, B, C). Likewise, hierarchical clustering analysis and PCA on the expression pattern of the myogenic and cardiomyogenic genes also show passage-dependent categorization (Figure 4D and 4E, Supplemental Table 1). After the induction, the mesenchymal stromal cells started to form multinucleated myotubes (Figure 4F).

**Figure 4.**
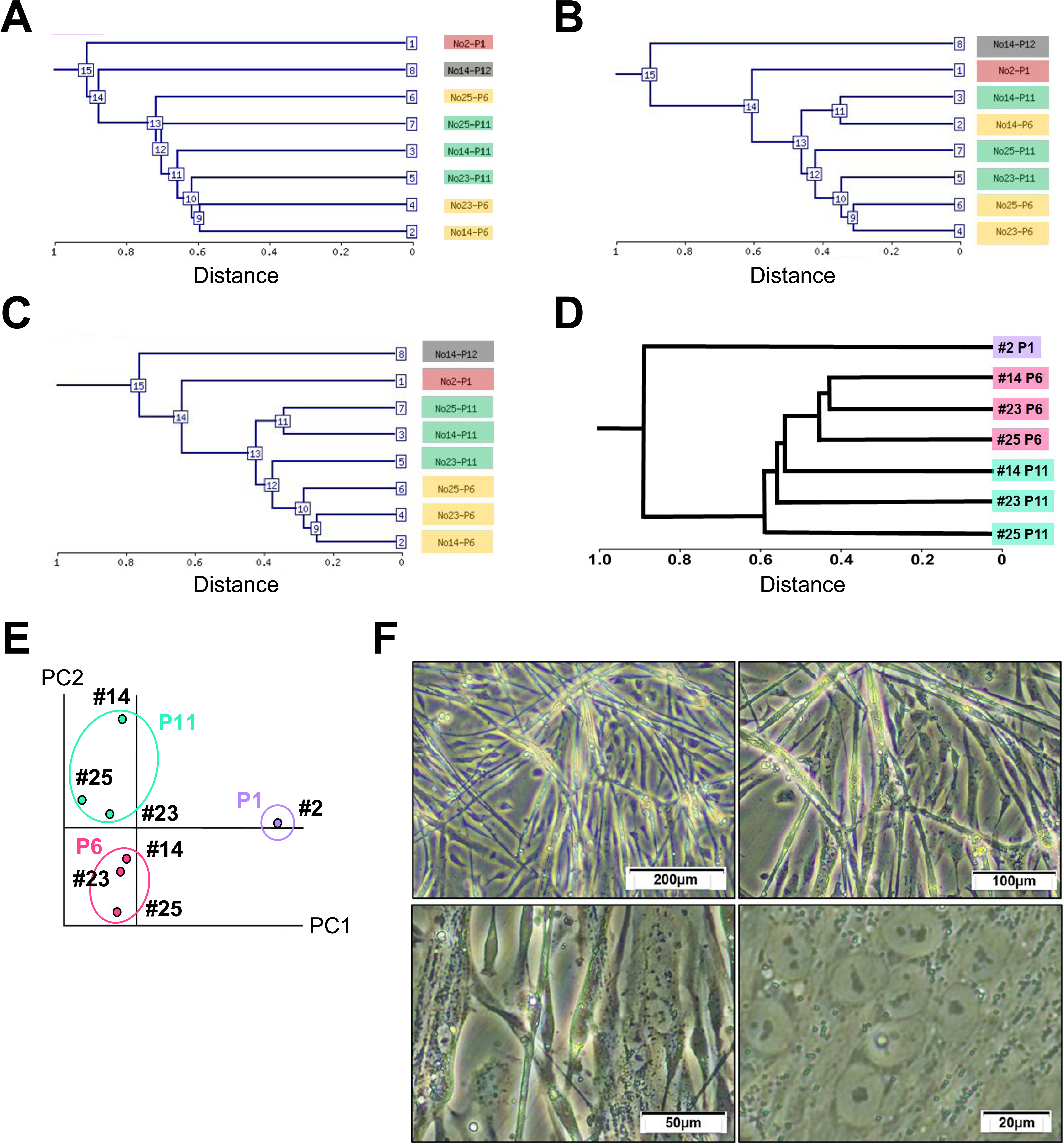
Global gene expression analysis of ESC-derived mesenchymal stromal cells. A. Hierarchical clustering analysis based on the expression of all genes (58,201 probes on Agilent SurePrint G3 Human GE v3 8x60K Microarray). B. Hierarchical clustering analysis based on expression levels of the mesenchyme-associated genes. C. Hierarchical clustering analysis based on expression levels of the stem cell-associated genes. D. Hierarchical clustering analysis based on expression levels of the muscle-associated genes. E. Principal component analysis of the muscle-associated genes. F. Phase-contrast photomicrographs of knst myogenesis.

## DISCUSSION

For the development of cell-based therapeutic strategies to genetic myogenic disorders, immortal cells as a raw material are required to gain sufficient number of cells, and detailed studies are therefore essential with regard to the characteristics of differentiated mesenchymal stromal cells. This present study demonstrated the detailed alterations of the mesenchymal stromal cells during expansion from P0 to P11 in monolayer culture. The fate of mesenchymal stromal cells generated from ESCs depended on passage number or population doubling levels in culture. In our previous study, we showed that human marrow stromal cells and umbilical cord blood-derived cells reach senescence, exhibit large, flat morphology at late passages and have different characteristics, depending on passage number and population doublings (Mori et al., 2005; Terai et al., 2005). Myogenic ability of ESC-derived mesenchymal stromal cells is possibly associated with surface markers, morphology, cytokines and differentiation capacity. c-kit, CD34, and CD140 serve as good markers to distinguish murine mesenchymal cells with multipotency, i.e. mesenchymal stem cells (Gojo and Umezawa, 2003). CD29+, CD44+, CD59+ and CD90+ cells from menstrual blood are capable of differentiating into myoblasts/myocytes and conferring human dystrophin expression in the murine model for Duchenne muscular dystrophy (Cui et al., 2007). In this study, we generated high-purity mesenchymal stromal cells for future myoblast therapy from a working cell bank of ESCs. The ESC-derived CD105+ cells with *in vitro* extensive proliferation capability exhibited myogenesis and genetic stability *in vitro*, implying that ESC-derived CD105+ cells are another cell sources for myoblasts in cell-based therapy to patients with genetic muscular disorders. Since ESCs are immortal, mesenchymal stromal cells generated from ESCs can be manufactured in a large scale with one lot in pharmaceutical purpose.

Mesenchymal stromal cells derived from ESCs have been examined from the viewpoints of differentiation propensity, surface markers, proliferation, and morphology (Barberi et al., 2005; Fu et al., 2015; Kimbrel et al., 2014; Moslem et al., 2015; Phillips et al., 2014; Trivedi and Hematti, 2008). They exhibit multipotency, i.e. adipogenic, osteogenic and chondrogenic differentiation *in vitro* (Barberi et al., 2005). They also show myogenic differentiation in vitro like mesenchymal stem cells derived from bone marrow, menstrual blood and placenta (Cui et al., 2007; Kawamichi et al., 2010; Kohyama et al., 2001; Mori et al., 2005; Okamoto et al., 2007; Umezawa et al., 1992). These mesenchymal stromal cells can be used for therapeutic agents or delivery vehicles to patients with graft versus host disease, ischemic heart disease, and lysosomal storage disorders. With the robust scalable manufacturing process described in this study, ESC-derived CD105+ cells serve as a starting material of these possible cellular therapy agents. ESC-derived CD105+ cells were mortal while the original ESCs (sees2) were immortal. The cells are, therefore, non-tumorigenic because they reach senescence or stop dividing after a limited number of replications. This limited cell lifespan could be an advantage from the viewpoint of tumorigenicity, but a disadvantage for scalable manufacturing of cell therapy products. iPSC-derived mesenchymal stromal cells exhibit almost the same phenotypes in differentiation propensity, surface markers, proliferation, and morphology as ESC-derived CD105+ cells (Chen et al., 2012; Phillips et al., 2014). Taken together, there is no great distinction in quality attributes of mesenchymal stromal cells derived from ESCs, iPSCs, and various tissues.

Implantation of myoblasts induced from ESC-derived mesenchymal stromal cells into patients with genetic muscular disorders is indeed an ideal strategy, from the viewpoint of industry-based, sustainable supply of large quantities of affordable, quality-controlled cells. It is unlikely that it is possible to prepare unaffected somatic cells in sufficient quantity, necessitating the use of stem cells from suitable, cost-effective allogeneic sources, such as ESCs and iPSCs. The cellular therapy products manufactured from ESCs and iPSCs can cover whole-body muscle because of their immortality. In addition, the proliferation capability and genetic stability of the ESC-derived mesenchymal stromal cells open up significant new possibilities in regenerative medicine. ESCs can be a promising cellular source for cell-based therapy to treat Duchenne muscular dystrophy, a lethal human disease for which no effective treatment currently exists (Assereto et al., 2016; De Paepe et al., 2016).

## MATERIALS AND METHODS

### Ethical statement

Human cells in this study were performed in full compliance with the Ethical Guidelines for Clinical Studies. The cultivation of hESC lines were performed in full compliance with “the Guidelines for Derivation and Distribution of Human Embryonic Stem Cells (Notification of the Ministry of Education, Culture, Sports, Science, and Technology in Japan (MEXT)) and “the Guidelines for Utilization of Human Embryonic Stem Cells (Notification of MEXT)”. The experimental procedures were approved by the Institutional Review Board (IRB) at National Center for Child Health and Development. Animal experiments were performed according to protocols approved by the Institutional Animal Care and Use Committee of the National Research Institute for Child Health and Development. All experiments with mice were subject to the 3 R consideration (refine, reduce, replace) and all efforts were made to minimize animal suffering, and to reduce the number of animals used.

### hESC culture

sees2 and sees5 were routinely cultured onto a feeder layer of freshly plated gamma-irradiated mouse embryonic fibroblasts (MEFs), isolated from ICR embryos at 12.5 gestations and passages 2 times before irradiation (30 Gy), in the hESC culture media. The hESC media consisted of Knockout^TM^-Dulbecco’s modified Eagle’s medium (KO-DMEM) (Life Technologies, CA, USA; #10829-018) supplemented with 20% 35 kGy-irradiated Knockout^TM^-Serum Replacement (KO-SR; #10828-028), 2 mM Glutamax-I (#35050-079), 0.1 mM non-essential amino acids (NEAA; #11140-076), 50 U/ml penicillin-50 μg/ml streptomycin (Pen-Strep) (#15070-063), and recombinant human full-length bFGF (Kaken Pharmaceutical Co., Ltd.) at 50 ng/ml. Cells were expanded using enzymatic passaging by recombinant trypsin (Roche Diagnostics, Indianapolis, USA).

### Manufacturing procedure

To generate embryoid bodies (EBs), sees2 and sees5 (5 × 10^^^3/well) were dissociated into single cells with 0.5 mM EDTA (Life Technologies) after exposure to the rock inhibitor (Y-27632: A11105-01, Wako, Japan), and cultivated in the 96-well plates (Thermo Fisher Scientific) in the EB medium [76% Knockout DMEM, 20% 35 kGy-irradiated Xeno-free Knockout Serum Replacement (XF-KSR, Life Technologies, CA, USA), 2 mM GlutaMAX-I, 0.1 mM NEAA, Pen-Strep, and 50 μg/mL l-ascorbic acid 2-phosphate (Sigma-Aldrich, St. Louis, MO, USA)] for 4 days. The EBs were transferred to T25 flasks coated with NMPcollagenPS (Nippon Meat Packers. Inc.), and cultivated in the XF32 medium [85% Knockout DMEM, 15% 35 kGy-irradiated XF-KSR, 2 mM GlutaMAX-I, 0.1 mM NEAA, Pen-Strep, 50 μg/mL l-ascorbic acid 2-phosphate, 10 ng/mL heregulin-1β (recombinant human NRG-beta 1/HRG-beta 1 EGF domain; Wako, Japan), 200 ng/mL recombinant human IGF-1 (LONG R3-IGF-1; Sigma-Aldrich), and 20 ng/mL human bFGF (Kaken Pharmaceutical Co., Ltd.)] for 60 to 70 days. The flasks were gently shaken to detach the cells. The detached cells were aggregated and could thus be easily removed by a pipette. The remaining adherent cells in the flasks were used for a resource of mesenchymal stromal cells. The adherent cells were then propagated in α-MEM medium supplemented with 10% FBS (Gibco or Hyclone) and 1% Pen-Strep for further in vitro and *in vivo* analysis.

### Karyotypic analysis

Karyotypic analysis was contracted out to Nihon Gene Research Laboratories Inc. (Sendai, Japan). Metaphase spreads were prepared from cells treated with 100 ng/mL of Colcemid (Karyo Max, Gibco Co. BRL) for 6 h. The cells were fixed with methanol:glacial acetic acid (2:5) three times, and placed onto glass slides (Nihon Gene Research Laboratories Inc.). Chromosome spreads were Giemsa banded and photographed. A minimum of 10 metaphase spreads were analyzed for each sample, and karyotyped using a chromosome imaging analyzer system (Applied Spectral Imaging, Carlsbad, CA).

### Gene chip analysis

Total RNA was extracted using TRIzol reagent (Thermo Fisher Scientific Inc.) according to the manufacturer’s instructions. RNA quantity and quality were determined using a Nanodrop ND-1000 spectrophotometer (Thermo Fisher Scientific Inc.) and an Agilent Bioanalyzer (Agilent Technologies, Santa Clara, CA). Total RNA was amplified and labeled with Cyanine 3 (Cy3) using Agilent Low Input Quick Amp Labeling Kit, one-color (Agilent Technologies) following the manufacturer's instructions. Briefly, total RNA was reversed transcribed to double-strand cDNA using a poly dT-T7 promoter primer. Primer, template RNA and quality-control transcripts of known concentration and quality were first denatured at 65°C for 10 min and incubated for 2 hours at 40°C with 5X first strand Buffer, 0.1 M DTT, 10 mM dNTP mix, and AffinityScript RNase Block Mix. The AffinityScript enzyme was inactivated at 70°C for 15 min. cDNA products were then used as templates for *in vitro* transcription to generate fluorescent cRNA. cDNA products were mixed with a transcription master mix in the presence of T7 RNA polymerase and Cy3 labeled-CTP and incubated at 40°C for 2 hours.

Labeled cRNAs were purified using QIAGEN’s RNeasy mini spin columns and eluted in 30 μl of nuclease-free water. After amplification and labeling, cRNA quantity and cyanine incorporation were determined using a Nanodrop ND-1000 spectrophotometer and an Agilent Bioanalyzer. For each hybridization, 0.60 μg of Cy3 labeled cRNA were fragmented, and hybridized at 65°C for 17 hours to an Agilent SurePrint G3 Human GE v3 8x60K Microarray. After washing, microarrays were scanned using an Agilent DNA microarray scanner.

Intensity values of each scanned feature were quantified using Agilent feature extraction software version 11.5.1.1, which performs background subtractions. We only used features which were flagged as no errors (Detected flags) and excluded features which were not positive, not significant, not uniform, not above background, saturated, and population outliers (Not Detected and Compromised flags). Normalization was performed using Agilent GeneSpring software version 13.0 (per chip:normalization to 75 percentile shift). There are total of 58,201 probes on Agilent SurePrint G3 Human GE v3 8x60K Microarray without control probes. Hierarchical clustering analysis and Principal Component Analysis were performed using NIA Array Analysis (https://lgsun.grc.nia.nih.gov/ANOVA/).

### Funding information

This research was supported by grants from the Ministry of Education, Culture, Sports, Science, and Technology (MEXT) of Japan; by Ministry of Health, Labor and Welfare (MHLW) Sciences research grants; by a Research Grant on Health Science focusing on Drug Innovation from the Japan Health Science Foundation; by the program for the promotion of Fundamental Studies in Health Science of the Pharmaceuticals and Medical Devices Agency; by the Grant of National Center for Child Health and Development. Computation time was provided by the computer cluster HA8000/RS210 at the Center for Regenerative Medicine, National Research Institute for Child Health and Development. We expand our acknowledgement to deanship of scientific research at King Saud University for funding this research through international research program "Metagenomics". AU thanks King Saud University, Riyadh, Kingdom of Saudi Arabia, for the Visiting Professorship.

## ACKNOWLEDGEMENTS

We would like to express our sincere thanks to Y. Takahashi and H. Abe for providing expert technical assistance, K. Miyado and H. Akutsu for fruitful discussion, to C. Ketcham for English editing and proofreading, and to E. Suzuki and K. Saito for secretarial work.

## COMPETING FINANCIAL INTERESTS

The authors declare no competing financial interests.

## AUTHOR CONTRIBUTION STATEMENT

AU designed experiments. YA, MS, and MM performed experiments. AU, YA, and MS analyzed data. MT contributed reagents, materials and analysis tools. CN, AU, MT and YA discussed the data and manuscript. AU and MS wrote this manuscript.

